## Supplementary Figures for "RecombineX: a generalized computational framework for automatic high-throughput gamete genotyping and tetrad-based recombination analysis"

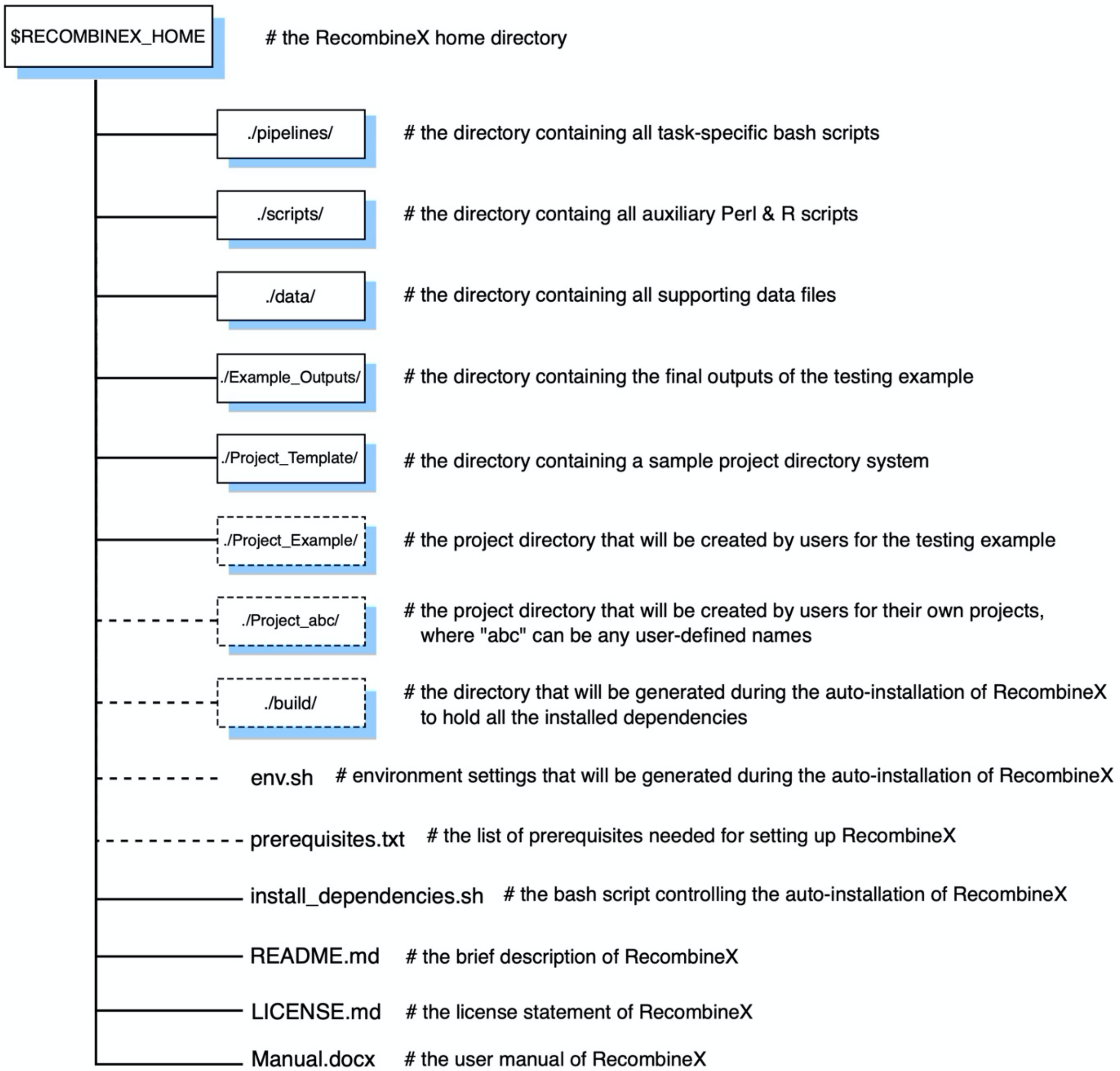


Figure S1. Overview of the RecombineX directory system. The pre-shipped top-level directories and individual files of RecombineX are denoted with solid lines. Additional directories and files to be generated during the installation of RecombineX are denoted with dashed lines.

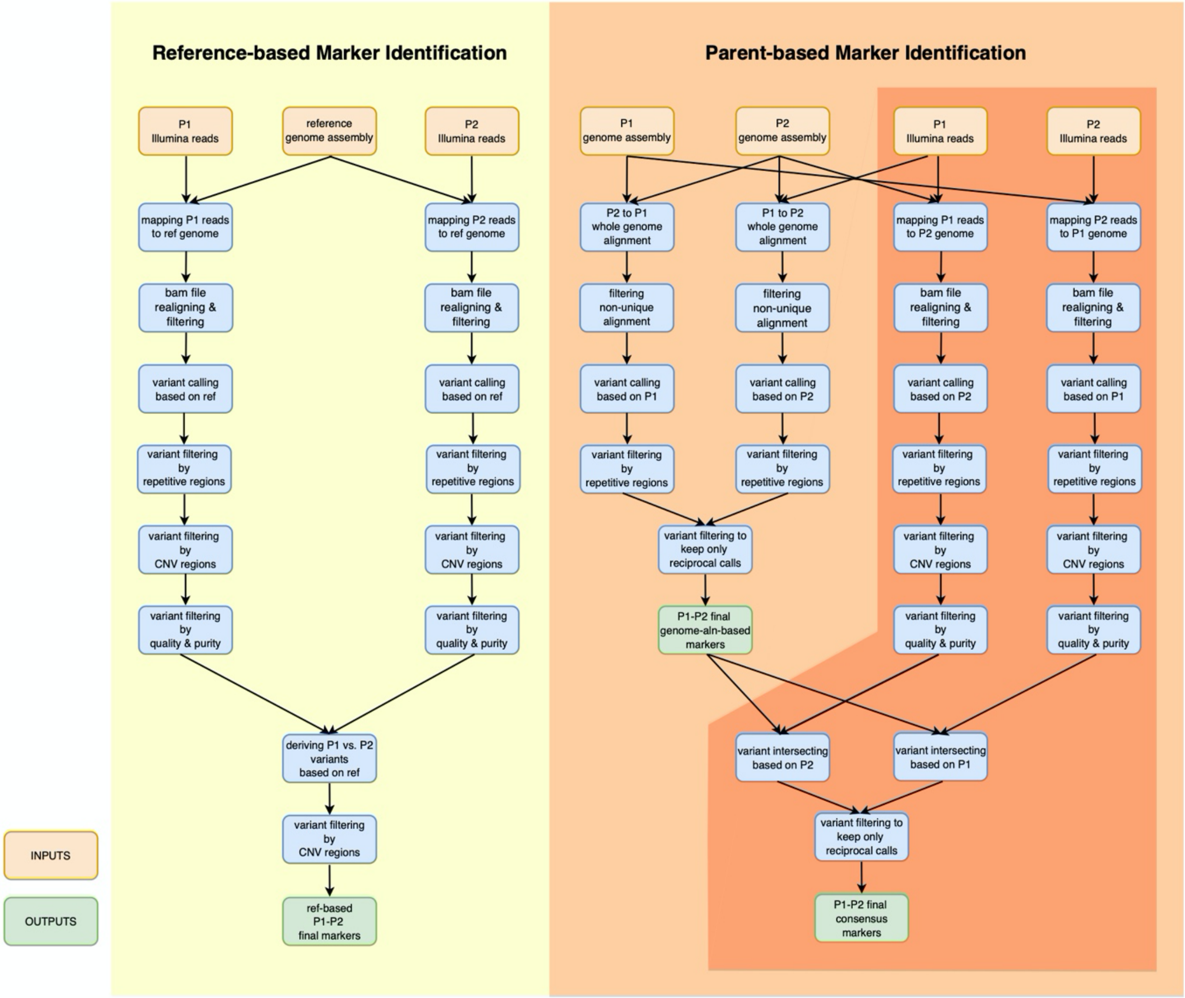


Figure S2. Overview of the RecombineX parental marker identification algorithms. Two marker identification modes are supported: the reference-based mode (colored in yellow) and the parent-based mode (colored in orange).

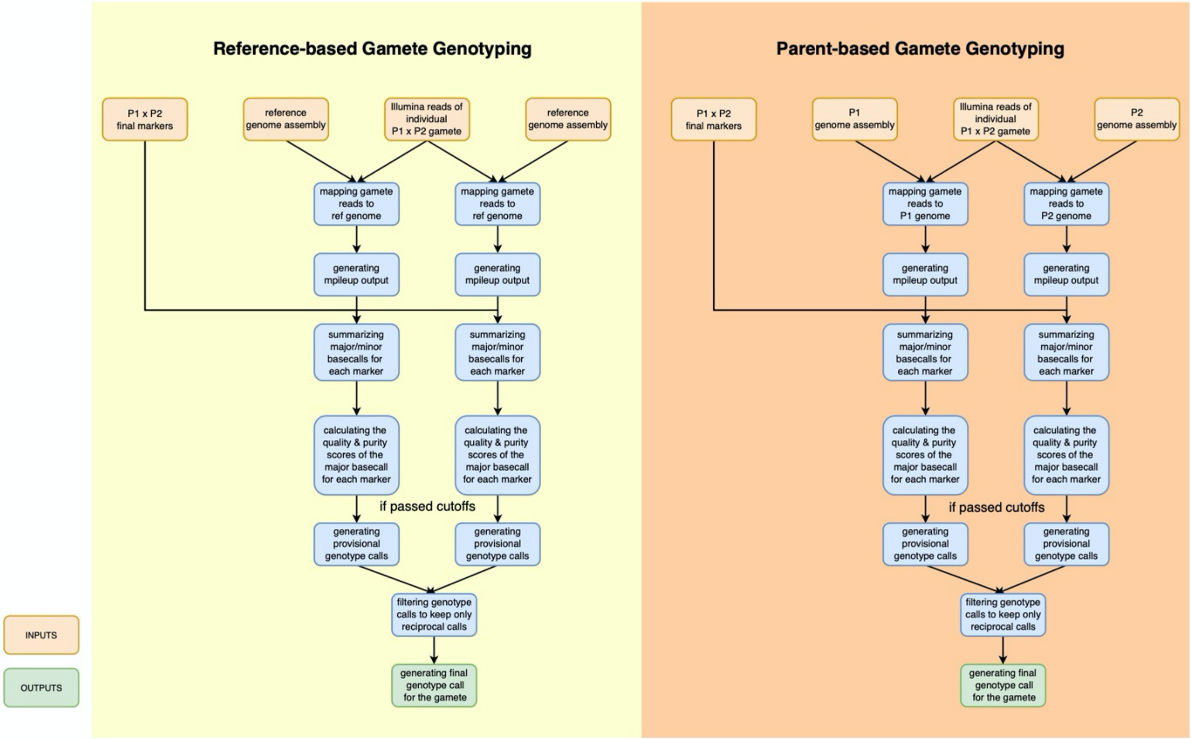


Figure S3. Overview of the RecombineX gamete genotyping algorithms. Two genotyping modes are supported: the reference-based mode (colored in yellow) and the parent-based mode (colored in orange).

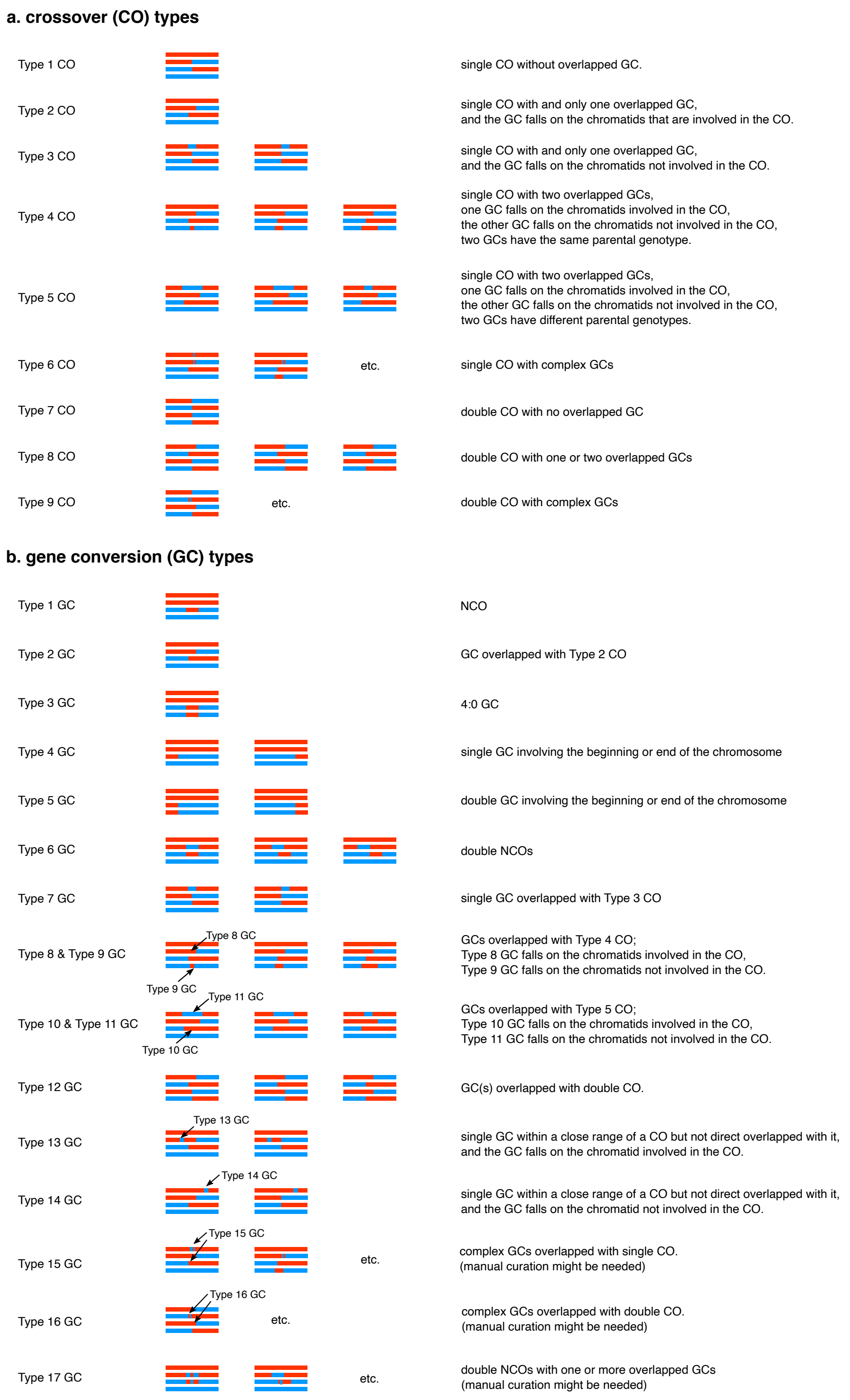


Figure S4. Overview of the RecombineX recombination event classification scheme. The definition and example of different CO and GC types are shown in panel a and panel b respectively. This classification scheme is designed based on the original ReCombine recombination event classification scheme (Anderson et al. PLoS One, 2011) with additional modifications.

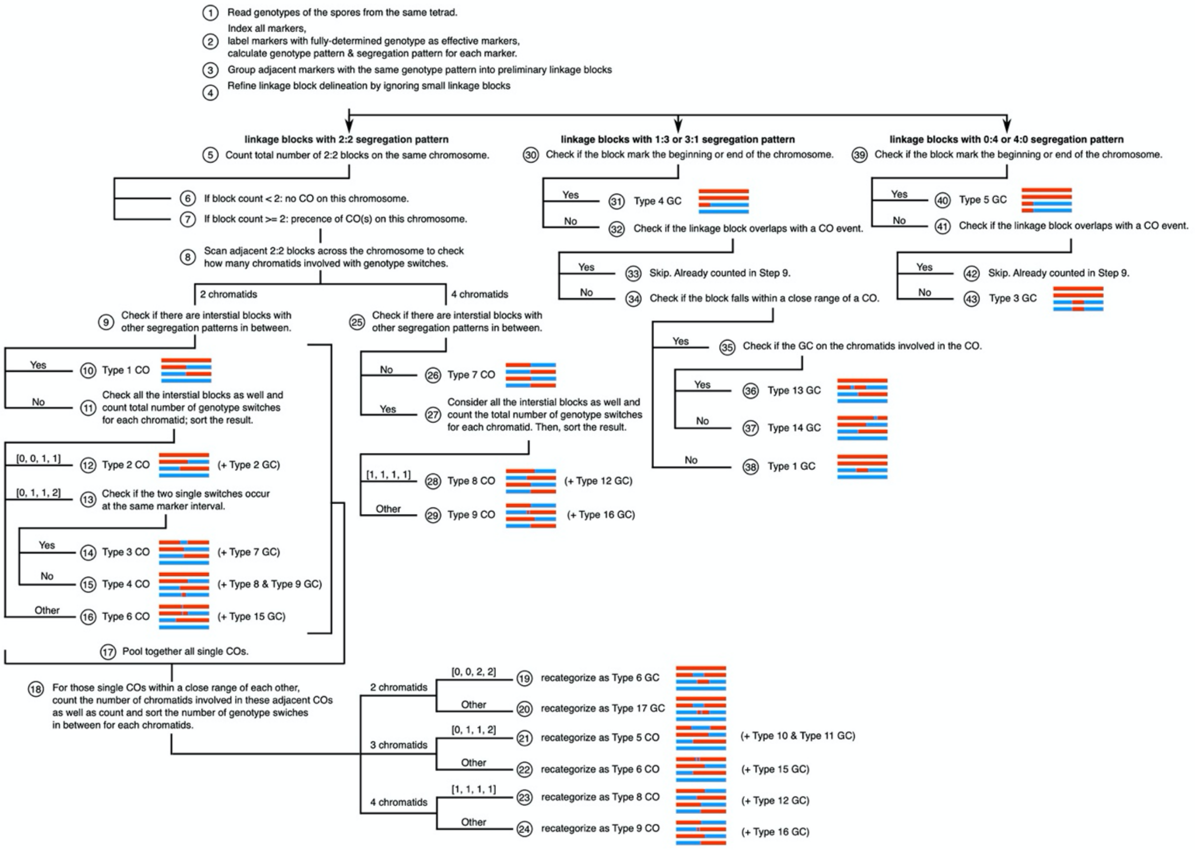


Figure S5. Overview of the RecombineX recombination event identification algorithm. This algorithm is designed based on the original ReCombine algorithm (Anderson et al. PLoS One, 2011) with additional modifications.

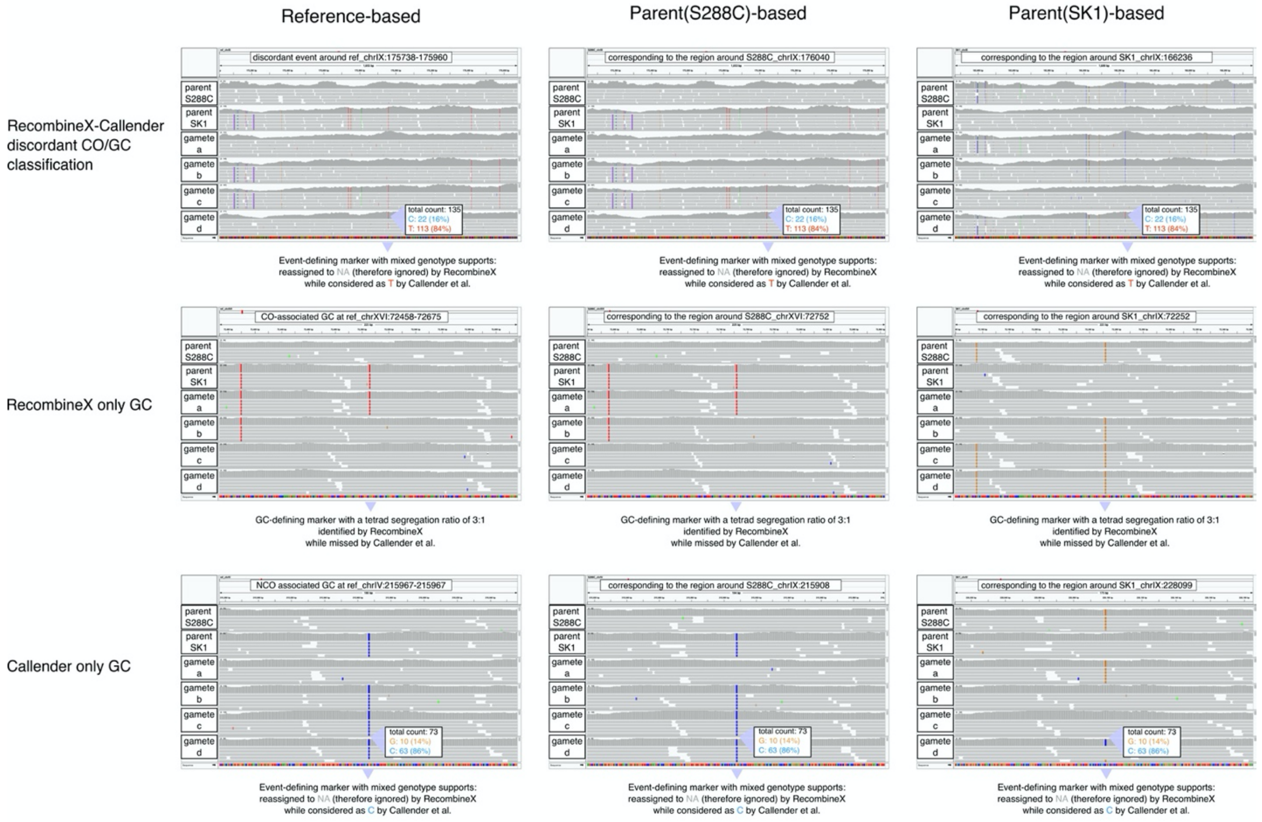


Figure S6. Manual examination of unmatched recombination events from the yeast S288C-SK1 tetrads in IGV. The read alignments of parent and gamete reads are visualized in IGV with event-defining SNP markers shown in colors.

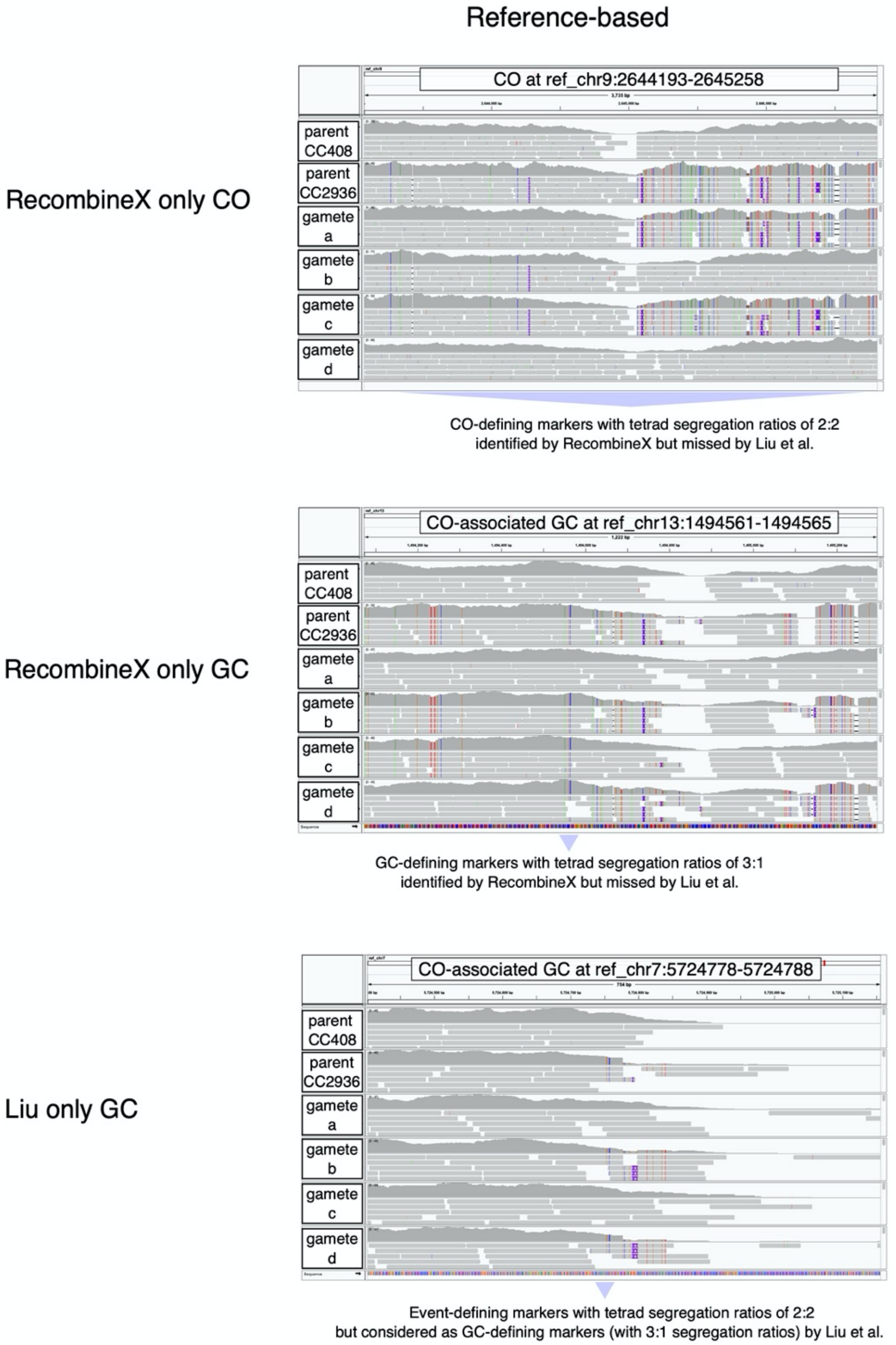


Figure S7. Manual examination of unmatched recombination events from the green alga CC408-CC2936 tetrads in IGV. The read alignments of parent and gamete reads are visualized in IGV with event-defining SNP markers shown in colors.
